# An ancient FMRFamide-related peptide-receptor pair induces defense behavior in a brachiopod larva

**DOI:** 10.1101/122085

**Authors:** Daniel Thiel, Philipp Bauknecht, Gáspár Jékely, Andreas Hejnol

## Abstract

Animals show different behaviors that can consist of various spatially or temporally separated sub-reactions. Even less complex organisms, like ciliated larvae that display important behaviors (e.g. metamorphosis, defense, feeding), need to coordinate coherent sub-reactions with their simple nervous system. These behaviors can be triggered by neuropeptides, which are short signaling peptides. Despite the high diversity of neuropeptides in animals, and although their immunoreactivity is widely used in morphological studies of animal nervous systems (e.g. FMRFamide), their function and role in trochozoan larval behavior has so far only been tested in a few cases. When mechanically disturbed, the planktonic larvae of the brachiopod *Terebratalia transversa* protrude their stiff and pointy chaetae in a defensive manner and sink down slowly: a startle reaction that is known from different chaetous trochozoan larvae. We found that both of these reactions can be induced simultaneously by the FMRFamide-related neuropeptide FLRFamide. We deorphanized the *Terebratalia* FLRFamide receptor and found its expression spatially separated in the apical lobe at the prototroch of the larvae and in the trunk musculature, which correlates with the tissues that are responsible to perform the two sub-reactions. A behavioral assay showed a decreasing efficiency of modified peptides in triggering this behavior, which correlates with the decreasing efficiency of activating the FLRFamide receptor in transfected CHO-K1 cells. Immunohistochemistry and *in situ* hybridization show FLRFamidergic neurons in the apical lobe as well as next to the trunk musculature. Our results show that the single neuropeptide FLRFamide can specifically induce the two coherent sub-reactions of the *T. transversa* startle behavior.

## Introduction

Planktonic organisms evolved different strategies to defend themselves from predation [1-4]. Morphological characters like shells, spines or chaetae [5-7] and behaviors such as vertical migration, contraction, active fleeing or passive sinking [8-11] can help to cope with certain predators. This is especially true for ciliated larvae that do not possess an elaborated nervous system and face the challenge of remaining in the water column for dispersal while avoiding predation. The startle behavior of several planktonic annelid and brachiopod larvae has often been described as a major defense strategy, where they stop swimming and protrude long and pointy chaetae [12-16]. The co-occurring sub-reactions of spreading the chaetae and stopping swimming take place in spatially separated tissues: the internal trunk musculature and the ciliated apical edge, respectively. Both sub-reactions have to be coordinated within the framework of a larval nervous system. One mechanism to achieve coordination of different reactions could be the use of neuropeptides as signaling molecules. Neuropeptides are known to influence many behaviors and can be crucial in the regulation and coordination of spatially or temporally separated coherent sub-reactions. During insect ecdysis for example, the eclosion hormone and the ecdysis-triggering hormone act both as a form of master-regulator on different peripheral as well as central targets, and each coordinates several sub-reactions [17-20]. Another example is neuropeptide Y which stimulates appetitive as well as consummatory ingestive behavior in the Siberian hamster [21]. In *Platynereis dumerilii*, myoinhibitory peptide first triggers settlement of the non-feeding larvae and later in development stimulates food intake in juveniles [22, 23].

Only a few studies have demonstrated the influence of neuropeptides on the behavior of trochozoan larvae: these studies show that neuropeptides can trigger settlement and influence their ciliary based locomotion [22, 24-26]. One of the neuropeptides that has been shown to influence ciliary beating of different trochozoan larvae is FMRFamide [24-26]. FMRFamide-immunoreactivity is widely used as a marker for neural substructures in morphological studies [27, 28]. Furthermore, while FMRFamide related peptides (FaRPs) have been identified in many metazoans, their phylogenetic relationship is difficult to reconstruct [29-31]. For the comparison of larval nervous systems it is therefore of crucial interest to understand the functional role of a neuropeptide and its versatility to trigger larval behaviors.

While experimental studies in trochozoan larvae are limited, the physiological effect of FMRFamide has been intensively investigated in adult trochozoans were it showed to have various muscular effects [32-36]. Depending on the species, it can increase or decrease the heartbeat [32], cause contractions or relaxation of somatic muscles [33-35] or change the efficiency of classical neurotransmitters on somatic muscles [37, 38]. So far only one study has shown a muscular effect of FMRFamide in larvae, which describes a twitching of the ciliated velum of *Tritia obsoleta* larvae [24]. Many immunohistochemical analyses on trochozoan larvae show FMRFamide-like immuno-reactivity associated with muscles or ciliary bands [39-41] but experimental data are mostly missing.

Since neuropeptides can act over longer distances [42], the localization of the neuropeptide receptor provides more information about the actually effected tissues than the peptide secreting cells that are labeled with the peptide antibodies. The majority of neuropeptide receptors are G-protein coupled receptors (GPCRs), with a few exceptions like insulin receptors or peptide gated ion channels [30, 43, 44]. For FMRFamide, there have been three different receptors deorphanized in invertebrates so far. One is an FMRFamide-gated amiloride-sensitive Na+ channel (FaNaCh) that has been identified in molluscs [45-47]. The two other receptors belong to two different groups of neuropeptide GPCRs. One of these FMRFamide-GPCRs was identified in the fruit fly *Drosophila melanogaster* [48, 49] and the other one in the annelid *P. dumerilii* [44]. This stands in contrast to many cases in which homologues ligands activate homologues receptors [44, 50, 51].

Here we show that the endogenous FaRP FLRFamide induces the characteristic defense behavior in the larvae of the brachiopod *Terebratalia transversa*, which consists of a downward sinking and the protrusion of their chaetae. Behavioral experiments and receptor deorphanization in combination with immunohistochemistry and *in situ* hybridization show that both coherent sub-reactions are specifically triggered by a single peptide via an ancient FaRP receptor. Together our results show how a single neuropeptide can trigger two coherent reactions and integrate evolutionary novelties such as trochozoan chaetae [52] into the *T. transversa* larval defense behavior.

## Material and Methods

### Collection and rearing of Terebratalia transversa larvae

Adult *T. transversa* (Sowerby, 1846) were collected in January by dredging in approximately 50-100 meter depth close to University of Washington’s Friday Harbor Laboratories, San Juan Islands. Larvae were obtained according to [53] by artificial fertilization and kept at 8-10°C. We used early larvae (2 days post fertilization) before chaetal formation and late larvae (4-5 days post fertilization) with clearly developed mantle lobes and long chaetae. For immunohistochemistry and *in situ* hybridization, larvae were relaxed in 7.8% MgCl_2_-6H_2_O for 10-15 min, fixed in 4% methanol-free formaldehyde in seawater for 1h, subsequently washed in PBS + 0.1% Tween and transferred into 100% methanol for storage at -20°C.

### Bioinformatics

The previously published transcriptome of *T. transversa* (Accession: SRX1307070) was queried for peptide precursor and receptor candidates using BLAST. For reference sequences, we used published FMRFamide like peptide precursor sequences from NCBI and checked candidates for signal peptides, cleavage sites and amidation sites. As reference sequences for the peptide receptor we used previously published collections [30, 44] and included transcriptomes of *Xenoturbella bocki* (Accession: SRX1343818), *Nemertoderma westbladi* (Accession: SRX1343819), *Meara stichopi* (Accession: SRX1343814) and *Halicryptus spinulosus* (Accession: SRX1343820) for additional sequences. The candidates were compared using the software CLANS [54] with a p-value cutoff of 1e-70. Sequences that were strongly connected in the cluster map were aligned with Clustal X v.2.1 [55], unconserved stretches were deleted manually and the best fitting amino acid substitution matrix was determined with Modelgenerator v.0.85 [56]. The final phylogenetic analysis was calculated with PhyML v3.0 [57] with 500 bootstrap replicates and visualized with Figtree v1.4.3 (http://tree.bio.ed.ac.uk/software/figtree).

### Behavioral assay

We tested synthetic peptides (GenScript) that were predicted from the prepropetide sequence and compared the reaction with non-native modifications of those.

To determine the efficiency of the native and modified peptides we tested at which concentration larvae contracted and spread their chaetae. To get an estimation of the peptide concentration that was necessary to induce a complete contraction we exposed larvae to a concentration of 50 nmol/l of the respective peptide and increased the concentration stepwise until the larvae contracted or a concentration of 50 *μ*mol/l was reached as an upper cutoff. Larvae were considered fully contracted when their chaetae were spread in all directions (fig.1 C). In addition to this, they did not further increase in contraction following an increase in peptide concentration, or followed by the use of the most sensitive peptide. Using this approximate concentration, we then tested fixed concentrations to narrow down the sensitivity window. Each test was performed with 30-100 larvae.

**Fig.1:**
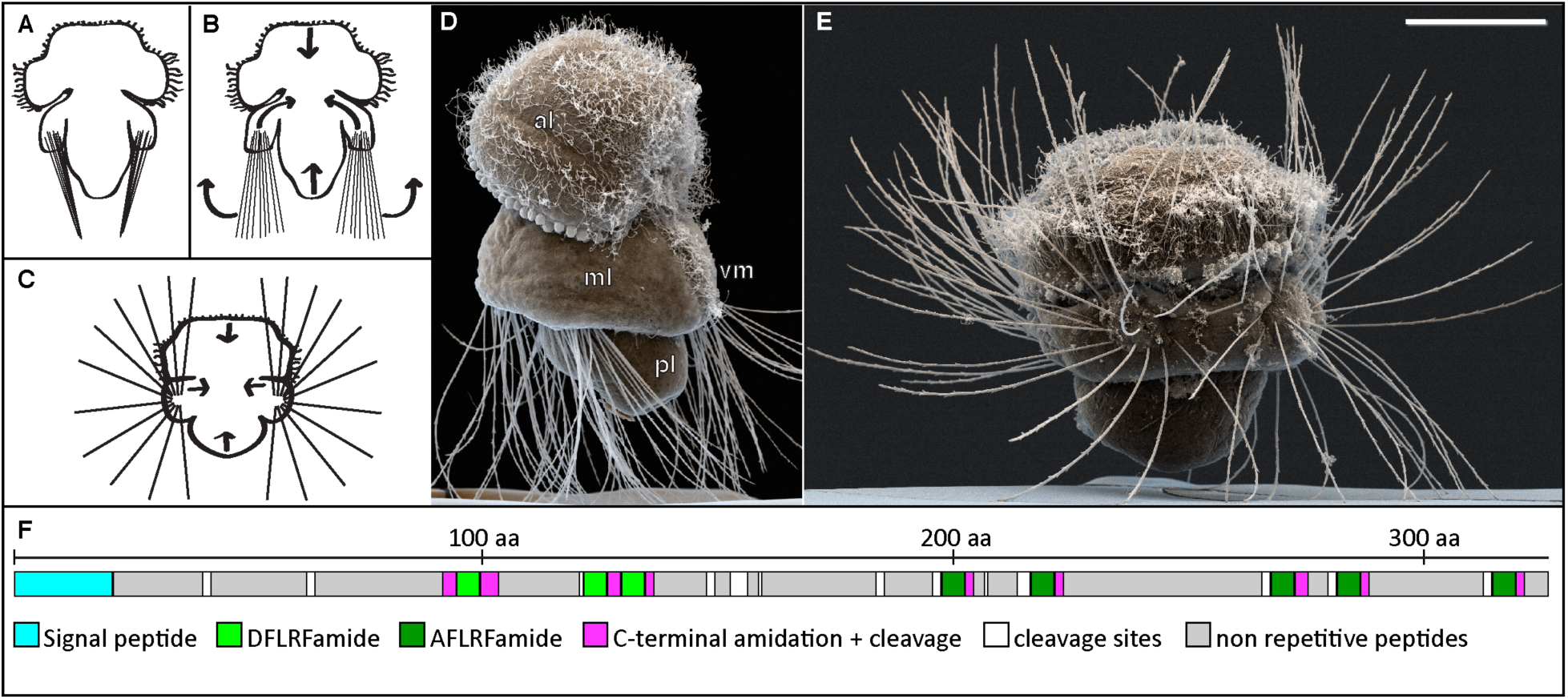
Defense reaction and FLRFamide prepropeptide of the *T. transversa* larvae. **A-C** sketch, **D-E** SEM photographs, anterior up, **F** sketch. **A** larva in relaxed stance during normal swimming; **B** non contracted larva that begins to spread its chaetae; **C** larva in defense stance with outspread chaetae; **D** non-contracted competent larva; **E** contracted competent larva with outspread chaetae. **F** schematic representation of *T. transversa* FLRFamide prepropeptide. al anterior lobe, ml mantle lobe, pl pedicle lobe, vm ciliated ventral mid line. Scale bar: 50 *μ*m

The vertical swimming experiments were recorded in 4.5 ml cuvettes with 50-100 larvae per cuvette, in a darkened box with top-illumination. About 1 minute after addition of the peptide in the treatments and water/DMSO in the controls we recorded the larval swimming (DMK 31AU03 camera, Imaging Source).

All experiments were repeated with at least one different batch of larvae from another fertilization and the outcome was averaged afterwards.

### Receptor deorphanization

For deorphanization we followed the procedure used by [44]: Full length ORFs of receptor candidate sequences were cloned into pcDNA3.1(+) mammalian expression vector (Sigma-Aldrich) and transfected into CHO-K1 cells together with a calcium sensitive luminescent apoaequorin-GFP fusion protein encoding plasmid (G5A) and a promiscuous Gα-16 protein encoding plasmid. After two days, coelenterazine h (Promega, Madison, WI, USA) was added and incubated with the cells for two hours. The luminescence response of the transfected cells was measured in a plate reader (BioTek Synergy Mx or Synergy H4, BioTek, Winooski, USA) over 45 seconds after addition of the neuropeptides. The response of the cells to 1 mM Histamine was used as a general control in each plate. All measurements for the dose response curves were made twice with different cell passages. Dose response curves were calculated using Prism 6 (GraphPad, La Jolla, USA) and normalized against the upper plateau values (100% activation).

### *In situ* hybridization

FLRFamide precursor and receptor sequences were amplified by PCR and cloned into pGEM Teasy vector (Promega) for *in vitro* transcription of DIG-UTP or DNP-UTP labeled RNA probes. For Tropomyosin we used a previously published clone [58]. The *in situ* hybridization protocol with an alternative hybridization buffer will be published elsewhere (Sinigaglia et al. 2017 - unpublished). In general we followed the protocol from [59] with following adjustments: Proteinase K treatment (10 *μ*g/ml) was adjusted to 8 minutes and the following postfixation was done in 3.7% Formaldehyde + 0.2% Glutaraldehyde in PBS + 0.1% Tween 20. The hybridization buffer contained 4 mol/l Urea, 5x SSC, 1% Dextran, 1% SDS, 50 *μ*g/ml Heparin, 50 *μ*g/ml single stranded DNA and no Formamide. The signal was developed with the TSA Plus Cy3 or Cy5 kit (Perkin Elmer) or NBT/BCIP as a substrate and detected via fluorescence or NBT/BCIP reflection [60] in a Leica SP5 confocal laser-scanning microscope.

### Immunohistochemistry

Customized polyclonal antibodies were raised in rabbits against CFLRFamide, coupled via a disulfide bridge to Keyhole limpet hemocyanin (GenScript^®^). Co-staining was either done with mouse anti-acetylated α-tubulin (Sigma, T6793) or mouse anti-actin (Seven Hills Bioreagents, LMAB-C4) antibodies. For immuno-staining we followed the protocol of [61] with the following adjustments: Proteinase K treatment (10 *μ*g/ml) was done for 3-5 minutes and after the proteinase inactivation step with glycine (2 mg/ml) the samples were incubated for 2-4h in PBS + 0.5% TritonX. Primary antibodies were incubated over 3 nights at 4°C and washed for 4-6 hours with at least 10 changes of washing medium. Secondary antibodies were incubated over night and included a secondary antibody without primary partner (goat anti rat) to test and subtract unspecific background staining. After washing the secondary antibodies for 4-6 hours with at least 10 changes of buffer, specimens were transferred into methanol and mounted in Murray’s clear (2:1 parts benzyl-benzoate: benzyl-alcohol).

## Results

### The endogenous neuropeptide FLRFamide triggers the defense behavior of *T. transversa* larvae

During normal swimming the chaetae of competent *T. transversa* larvae rest against their pedicle lobe with their tips forming a bundle (fig.1 A). When the larvae get disturbed (e.g. mechanical irritation with a pipette tip) they stop swimming, sink down slowly and exhibit a defensive stance by lengthwise contraction of their body to spread the 4 bundles of chaetae outwards (fig.1 B,C,E). At maximal contraction the larvae spread their chaetae likewise in all directions to surround their soft body (fig.1 C,E).

We discovered a FLRFamide prepropeptide sequence in the transcriptome of *T. transversa*. The FLRFamide precursor contains a signal peptide, three copies of DFLRFamide and five copies of AFLRFamide, partially separated by intermediate sequences (fig.1 F). When we exposed larvae to synthetic FLRFamide, they contracted lengthwise, spread out their chaetae and sank down slowly. Both predicted peptides, DFLRFamide and AFLRFamide, caused the same behavior. When exposed to 50 nmol/l DFLRFamide all larvae showed initial signs of contraction, indicated by their chaetae bundles being slightly fanned out while still pointing in a posterior direction (fig.1 B,D). About half of the larvae continued swimming while the other half started to sink slowly on the bottom. An increase in peptide concentrations lead to an overall increase in contraction, resulting in a stronger spreading of the chaetae and more larvae sinking down. A maximum contraction of all larvae, with their body being completely surrounded by chaetae was observed at concentrations of 500-750 nmol/l DFLRFamide. When we removed the peptides by exchanging the medium with fresh seawater, all larvae returned to normal swimming behavior. Continuous exposure for about 2 hours led to a desensitization and the larvae resumed normal swimming without removing the peptides. When the larvae were desensitized by continuous exposure to DFLRFamide, they became also insensitive to AFLRFamide and vice versa.

Taken together we found *T. transversa* larvae show a continuing behavior which is similar to their startle response when we exposed them to one of the neuropeptides encoded on the endogenously expressed FLRFamide precursor.

### FLRFamide causes sinking of larvae independent from the protrusion of their chaetae

One part of the defense behavior of *T. transversa* larvae is a slow downward sinking. This reaction can already be observed in early larvae (sFig.1 E) before they develop long chaetae. Due to the shape of the larvae and the lack of a clearly restricted prototroch it was not possible to directly record the ciliary beating. However, since their larval locomotion is purely driven by ciliary beating, we hypothesize that FLRFamide influences the ciliary movement. To measure the swimming behavior in an unbiased manner, we recorded the position of freely swimming larvae in vertical columns and compared it to the position of larvae after exposure to DFLRFamide (fig.2 A). To test the possibility that the sinking is caused by an increase in the water drag due to the protruded chaetae, we also recorded early larva that do not have long chaetae yet but already express FLRFamide in the apical lobe (sFig.1 D). Both stages showed a sinking behavior that shifted the distribution of the larvae in the water column down, compared to the controls (fig.2 B).

**Fig.2:**
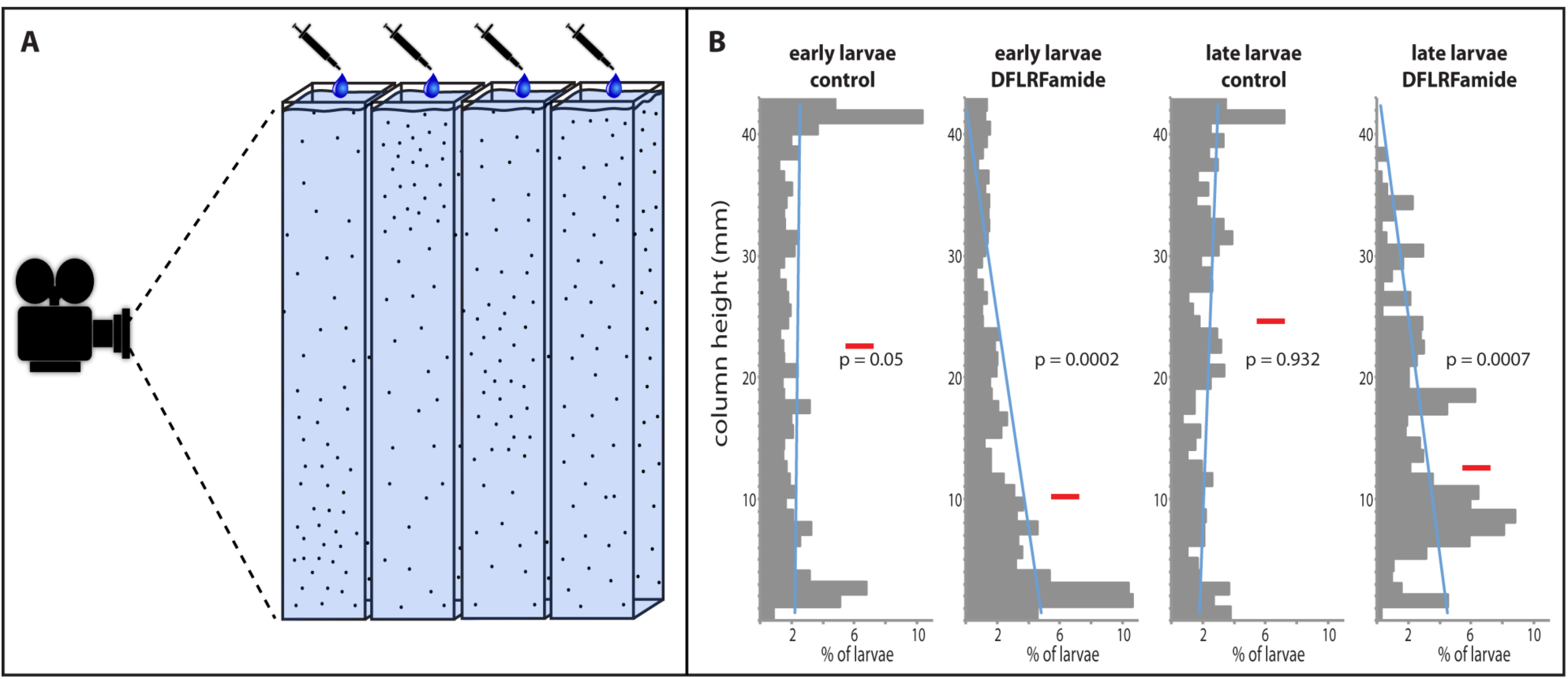
Influence of FLRFamide on the vertical distribution of early and late larvae. **A** Schematic experimental setup to compare the vertical distribution of larvae under different conditions in parallel. **B** Vertical distribution of early and late larvae in the water column and the effect of 5 *μ*mol/l DFLRFamide. Red bar shows average level of swimming height, p values are calculated for difference in distribution of larvae in upper versus lower half of the column (2-tailed, unpaired t-test), blue line is the estimated trend line (not statistically supported). Distribution was measured over a period of 5 seconds, about 1 minute after exposure to peptide.

### Modified peptides trigger the defense behavior at different concentration thresholds

We tested at which concentration modified peptides induce the contraction that leads to the erection of the chaetae, with 50 *μ*mol/l as a cutoff for the maximum concentration (tab. 1, sTab. 1). The larvae were most sensitive to DFLRFamide and showed full contraction (fig. 1 C,E), sinking and very slow movement on the bottom of the dish at concentrations between 500 nmol/l and 750 nmol/l (batch dependent). Further increasing the concentration did not lead to an obvious increase in the reaction. AFLRFamide showed to be slightly less effective by triggering full contraction of all larvae between 1 and 1.5 *μ*mol/l.

The reduced peptide sequence FLRFamide showed to be effective at 3 *μ*mol/l. Changing the amidated C-terminal phenylalanine to an amidated tryptophan reduced the effectiveness by about 10-fold, with a minimum necessary concentration of 7.5 *μ*mol/l of DFLRWamide or 20 *μ*mol/l of AFLRWamide. Changing the C-terminal phenylalanine to the non-aromatic leucine only led to a very weak contraction in some of the larvae at 50 *μ*mol/l DFLRLamide (fig.1 B) whereas 50 *μ*mol/l AFLRLamide gave no reaction at all. Reducing the sequence to the three C-terminal amino acids LRFamide also didn’t lead to any contraction; neither did any of the other non-related peptides that we tested. (The full list of tested peptides is given in sTab.01). The overall most effective versions were the ones that are encoded on the pro-peptide sequence, DFLRFamide and AFLRFamide. The reduced peptide FLRFamide was slightly less effective and a modification of the amidated C-terminus reduced the effectiveness even more.

### Identification of the *T. transversa* FaRP receptor

Based on BLAST e-value similarities and cluster analysis we tested four receptor candidates for their activation by FLRFamide (fig.3 A). One candidate belongs to a cluster of receptors that includes the deorphanized *P. dumerilii* FMRFamide receptor [44] with related sequences in all major bilaterian groups including Xenacoelomorpha (fig.3 A “I”). The second candidate belongs to the luqin receptors (fig.3 A “II”). The third candidate belongs to a group of related receptors with unknown ligand (fig.3 A “III”) and the fourth one belongs to a group that shows similarities with the deorphanized *Drosophila melanogaster* FMRFamide and the *P. dumerilii* NKY receptors (fig.3 A “IV”). Transcriptome searches for an FMRFamide gated ion channel (FaNaCh) as it was identified in molluscs did not reveal any orthologs in *T. transversa*, even when using FaNaCh orthologs that were identified in the brachiopods *Lingula anatina* and *Novocrania anomala*.

**Fig.3:**
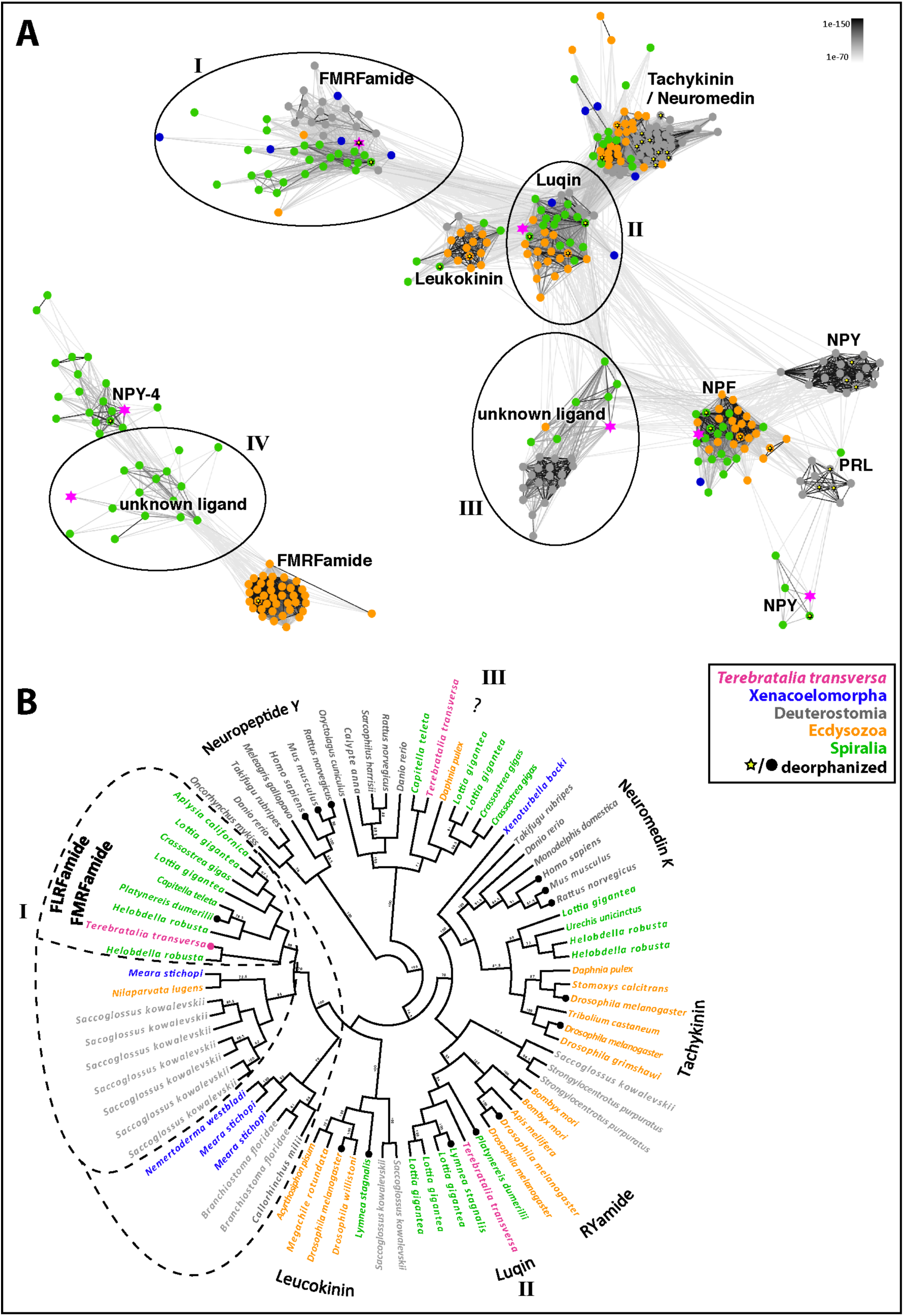
Phylogeny of the *T. transversa*-FLRFamide receptor. **A** Clustermap of metazoan neuropeptide GPCRs that show connections to FMRFamide receptors. Connections correspond to blastp connection with p-values <1e-70. Groups that include the receptors I-IV that were tested for activation by FLRFamide are encircled. NPY neuropeptide Y, NPF neuropeptide F, PRL prolactin releasing peptide. **B** Cladogram of neuropeptide GPCRs that showed connections in the clustermap to the *T. transversa* FLRFamide receptor (I). The dashed lines indicate receptor groups related to the *Terebratalia* FLRFamide receptor. Branches with filled circle at the end indicate a receptor that was deorphanized in a previous study.

To test whether FLRFamide is the ligand of one of these receptors, we tested their activation by DFLRFamide in transfected CHO-K1 cells. This first test showed that only the candidate that is related to the previously deorphanized *P. dumerilii* FMRFamide-receptor was activated by 1 *μ*mol/l DFLRFamide but none of the other candidates. We called this GPCR the FLRFamide receptor. Since DFLRFamide already triggered the defense stance at concentrations below 1 *μ*mol/l in the behavioral assay, we did not test the negative GPCR candidates at higher peptide doses. We further compared the luminescence response of FLRFamide receptor expressing CHO-K1 cells to 1 *μ*mol/l DFLRFamide, AFLRFamide, FLRFamide, DFLRWamide and DFLRLamide (fig.4 A). The two native forms DFLRFamide and AFLRFamide lead to the highest luminescence, followed by FLRFamide in a similar range. DFLRWamide gave a strongly decreased luminescence and the values of DFLRLamide were barely higher than the negative control.

**Fig.4:**
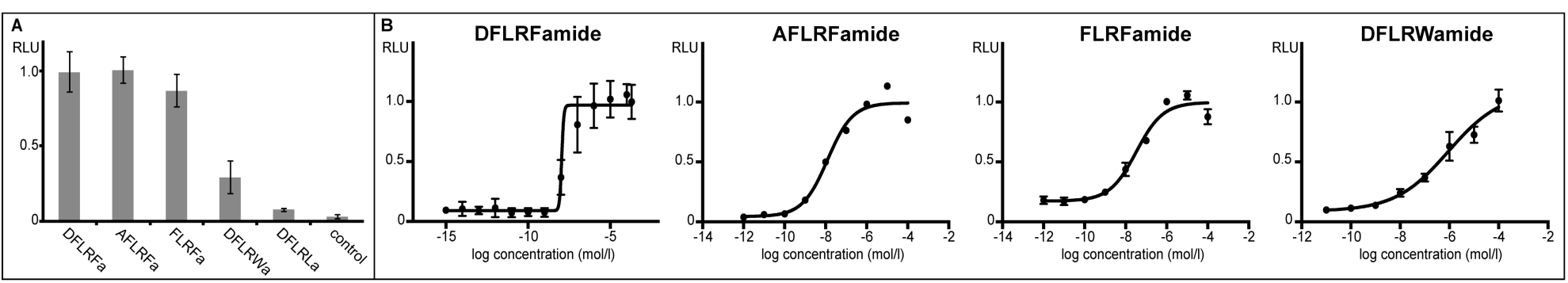
Luminescence response of *T. transversa*-FLRFamide receptor expressing CHO-K1 cells to different peptides. A Relative luminescence of *T. transversa*-FLRFamide receptor expressing cells after exposure to different peptides with a fixed concentration of 1 *μ*mol/l. B Dose response curves of *T. transversa*-FLRFamide receptor expressing cells to different concentrations of DFLRFamide, AFLRFamide, FLRFamide and DFLRWamide. Luminescence values are given relative to maximum luminescence (max = 1). RLU relative luminescence.

Dose response curves were recorded for DFLRFamide, AFLRFamide, FLRFamide and DFLRWamide (fig.4 B) and EC_50_ values (half maximal effective concentration) were determined for each peptide.

DFLRFamide and AFLRFamide showed EC_50_ values in a similar range (1.12E-08 and 1.24E-08 mol/l). The EC_50_ value for FLRFamide was about three times higher (3.32E-08 mol/l) and the one for DFLRWamide was the highest of all tested peptides (9.06E-07 mol/l). The EC_50_ values are listed in table 1, together with the concentrations that were necessary to trigger the defense stance in the behavioral assay.

**Tab.1:** Necessary peptide concentrations to evoke larval defense stance compared to EC_50_ values of receptor activation.

| peptide | necessary concentration to induce full contraction | $\text{EC}_{50}$ receptor assay |
| --- | --- | --- |
| **DFLRFamide** | 0.625 $\mu\text{mol/l}$ | 11.2 nmol/l |
| **AFLRFamide** | 1.5 $\mu\text{mol/l}$ | 12.4 nmol/l |
| **FLRFamide** | 3 $\mu\text{mol/l}$ | 33.2 nmol/l |
| **DFLRWamide** | 8.75 $\mu\text{mol/l}$ | 0.9 $\mu\text{mol/l}$ |

After we deorphanized the FLRFamide receptor, we tested its phylogenetic relationship to the receptors that showed connections in the cluster analysis (fig.3 A “I-III”). We did not include the unrelated *T. transversa* orphan receptor that is related to the insect FMRFamide and trochozoan NKY receptors (fig.3 A “IV”). The *T. transversa* FLRFamide receptor is directly related the previously deorphanized *P. dumerilii* FMRFamide receptor and several orphan receptors of other trochozoan (fig.3 B). Orthologs to these trochozoan FMRFamide/FLRFamide receptors were found in the insect *Nilaparvata lugens*, the hemichordate *Saccoglossus kowalevskii* and the xenacoelomorph *Meara stichopi*. A third related group includes orphan receptors from the cephalochordate *Branchiostoma floridae*, the vertebrate *Callorhinchus milii* and the xeacoelomorphs *M. stichopi* and *Nemertoderma westbladi*, These receptors form a fully supported group of neuropeptide GPCRs with homologs in all major bilaterian clades that is well seperated from other neuropeptide GPCR groups. (Fig.3 B)

Taken together we discovered that the *T. transversa* FLRFamide receptor belongs to an ancient neuropeptide receptor group and is efficiently activated by the two peptides AFLRFamide and DFLRFamide that are encoded on the *T. transversa* prepropeptide sequence.

### *In Situ* Hybridization and Immunohistochemistry show localization of peptide receptor in trunk musculature and apical prototroch region

The FLRFamide precursor has several expression domains within the apical lobe around the neuropile, and two domains on the ventral side at the anterior border of the mantle lobe (fig.5 A,B,E; sFig.1 B,C). The number of domains in the apical lobe varied between three and five (fig.5 A,B) and each domain consists of approximately three to seven cells. The combined *in situ* hybridization with *tropomyosin* as a marker for the musculature shows that the FLRFamide precursor expression in the mantle lobe is adjacent to the ventral side of the trunk musculature (fig.5 E). The FLRFamide receptor is expressed in a left and a right stripe in the trunk musculature (fig.5 B,C; sFig.1 A) as well as in the musculature that projects and surrounds the chaetae sacs (fig.5 D; sFig.1 A). Apart from the expression in the musculature the receptor is also expressed in the apical lobe in a broad stripe underneath the ciliated prototroch (fig.5 B,C; sFig.1 A).

**Fig.5:**
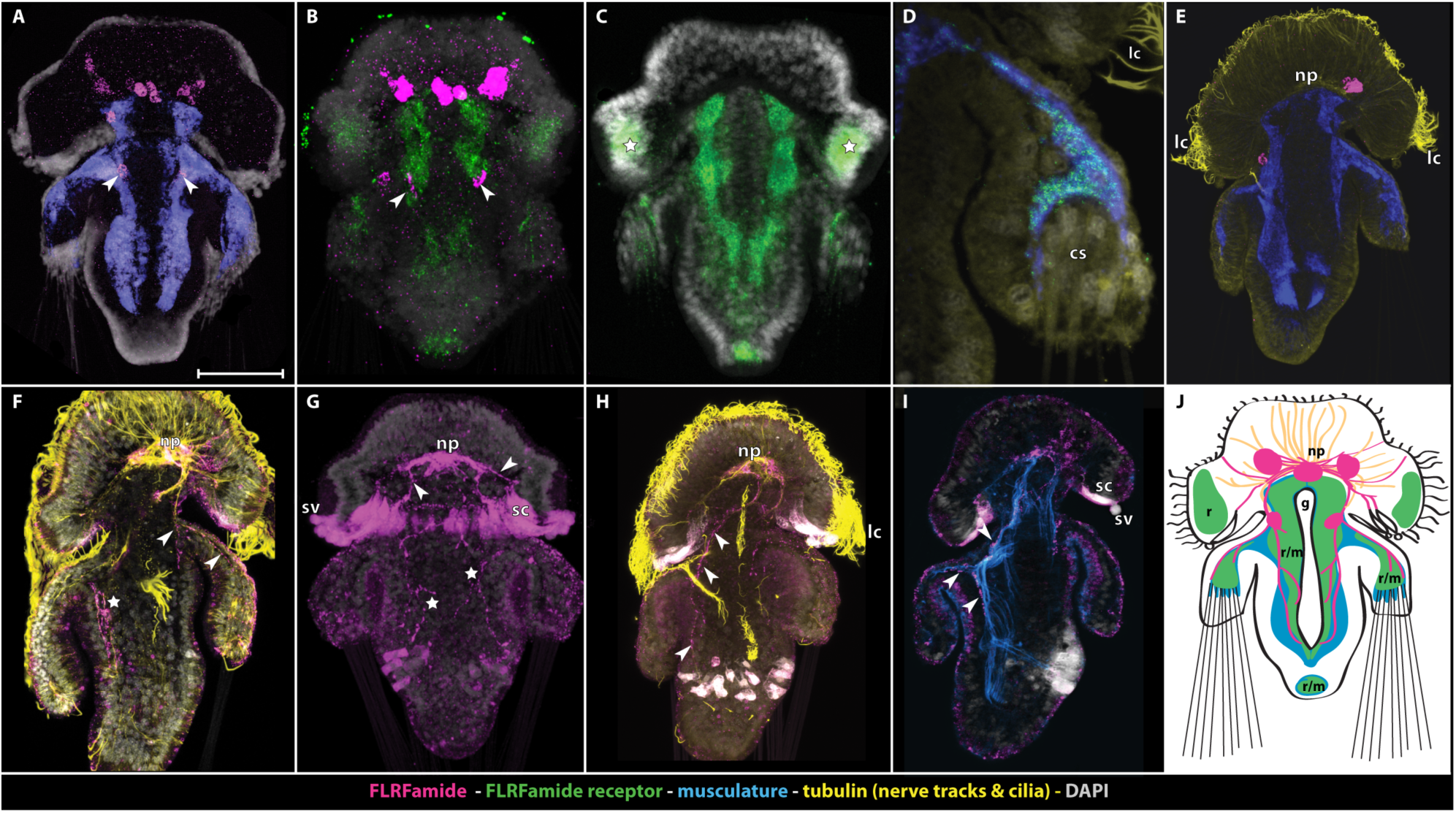
*In situ* hybridization and immunostainings of *Terebratalia*-FLRFamide, *Terebratalia*-FLRFamide receptor, musculature and tubulin. **A**-**E** *in situ* hybridization; **F**-**I** immunohistochemistry; **A-D,G,J** front view; **E,F,H,I** side view, ventral side left. **A** *FLRFa* and *tropomyosin* expression, arrows show *FLRFa* expression in mantle lobe. **B** *FLRFa* and *FLRFa receptor* expression, arrows show *FLRFa* expression in mantle lobe. **C** *FLRFa receptor* expression, stars show expression underneath prototroch. **D** *tropomyosin* and FLRFa receptor co-expression around chaetae sacks. **E** *FLRFa* and *tropomyosin* expression. **F** FLRFa and tubulin staining, star shows branching of FLRFa positive nerves inside ventral trunk area, arrows show branching of dorsal FLRFa positive trunk-nerve towards chaetae sacks. **G** FLRFa staining, stars show branching of FLRFa positive nerves inside dorsal trunk area, arrows show projections into secretory cells underneath the prototroch. **H** FLRFa and tubulin staining, arrows show nerve projecting from neuropil into ventral part of the trunk. **I** FLRFa and actin staining, arrows show lining of the musculature by FLRFamidergic nerves, projecting into mantle and posterior part of the trunk. **J** Schematic drawing of FLRFamidergic cells and nerves, FLRFa receptor and musculature. *cs* chaetae sacks, *g* gut, *lc* locomotory cilia of the prototroch, *m* musculature, *np* neuropil, *r* receptor, sc secretory cells, *sv* secretory vesicles. Color code is indicated at the bottom of the figure plate: Magenta FLRFamide (A,B,E precursor expression; F-I processed peptides), green FLRFamide-receptor, blue musculature (A,D,E tropomyosin expression; I actin staining), yellow nerve tracks and cilia.

Since *in situ* hybridization only reveals were the peptide precursor is expressed, we used antibody staining to visualize the nerves that secrete the active peptides. The customized FLRFamide antibody revealed immuno-reactive longitudinal nerves that project from the apical neuropil (fig.5 F,G,H,I) pairwise along the ventral (fig. 7 H) and dorsal (fig. 7F) side into the trunk after branching off into the mantle towards the chaetae sacks at the border of the apical lobe and the mantle lobe (fig.5 F,H,I)). The nerves in the trunk are branching of strongly on the ventral side (fig.5 F,G) and are at least partially directly adjacent to the musculature (fig.5 I). The neuropile shows generally strong FLRFamide immunoreactivity with some nerves projecting towards the apical ciliary band and into the secretory cells that continue into secretory vesicles outside the apical lobe underneath the prototroch (fig.5 G,H,I). The secretory cells and vesicles themselves are prone to antibody trapping so no statement can be made whether they in fact contain FLRFamide (compare fig.5 G without background subtraction and fig.5 F,H,I with background subtraction in secretory cells and secretory vesicles).

## Discussion

### FLRFamide triggers two coherent reactions via an ancient FaRP receptor

The receptor deorphanization and phylogenetic analysis shows that the *Terebratalia* FLRFamide receptor belongs to the ancient FaRP-GPCR group with closely related trochozoan GPCRs that include the deorphanized *P. dumerilii* FMRFamide receptor [44] and related orphan GPCRs in all major bilaterian groups (fig.3).The comparable sensitivity to different peptide modifications of the larvae in the behavioral assay and the EC_50_ values of the receptor cell assay suggests that the larval response is triggered via the FLRFamide receptor. The expression of the FLRFamide receptor in the longitudinal trunk musculature and the musculature adjacent to the chaetae sacks in the mantle supports a direct mode of signaling whereby FLRFamide directly triggers the protrusion of the chaetae by inducing a muscle contraction. The expression of the receptor in a broad stripe underneath the ciliary band and the sinking of early and late larva, independent of the presence or absence of chaetae, also supports a direct effect of FLRFamide on the ciliated cells to induce the sinking behavior. While a direct influence of FMRFamide on the ciliary movement of trochozoan larvae has already been suggested before [25, 26], the combination of this reaction with the muscular contraction observed in the *T. transversa* larval startle response consists of two different behavioral actions.

### The advantage of coherent sub-reactions during *T. transversa* defense behavior and their control by a single peptide

Neuropeptides are considered to be ancient signaling molecules that are used in complex as well as simple nervous systems and are even present in Placozoa that lack neurons entirely [30, 62]. There are a few examples of complex behaviors that involve coherent sub-reactions like insect ecdysis or feeding, which are known to deploy single neuropeptides to act on several targets as a form of master-regulator [17-21]. When a single neuropeptide controls the erection of chaetae and sinking, it might coordinate the startle reaction independent from a direct neuronal wiring between these two structures.

While many zooplankton organisms escape potential predators by a sudden increase in velocity, some species have been observed to use passive sinking as an efficient escape strategy instead [3, 63]. Passive sinking seems efficient for slow animals to escape quicker predators such as copepods that do not detect their prey by vision but by sensing water disturbance [3, 11, 64]. It has been described that other brachiopod and annelid larvae seem to have a similar startle behavior as *T. transversa* [12-16]. Direct observations showed that tiny fishes spit out *Sabellaria* larvae with their spines erected [15] and experimental data showed that *Sabellaria* larvae with long chaetae have a higher survival rate compared to younger larvae without chaetae when exposed to different predators [14]. The combination of a passive sinking behavior while actively erecting chaetae might thus increase the chance to escape different predators when compared to either only showing sinking behavior or a protrusion of chaetae.

Our results demonstrate a case in which a single receptor-ligand pair can trigger two coherent reactions that integrate evolutionary novelties such as trochozoan chaetae [52] and ancient traits such as ciliary based locomotion [65] into the *T. transversa* larval startle behavior.

### FaRP receptor-ligand pair was redeployed several times during trochozoan evolution

The conserved receptor ligand pair in *P. dumerilii* and *T. transversa* stands in contrast to the various actions that FMRFamide can have in trochozoans. Several studies on trochozoan larvae have shown that FMRFamide like immuno-reactive nerves can be associated with several structures in a single animal and often include a combination of the apical organ, ciliary bands and the musculature, which would suggest different regulatory roles [39-41, 66-68]. However, in this context it is important to mention that antibodies against FMRFamide strongly cross-react with other peptides ending in RFamide, even within the same specimen [69, 70]. Inter-species comparisons of such labeled neurons across larger evolutionary distances are thus problematic. The cross-reactivity of polyclonal FMRFamide antibodies has even led to the discovery of new peptides in the past [71, 72] and the C-terminal ending RFamide is part of many different neuropeptides [30, 70, 73, 74]. Only a few experimental studies exist on the effect of FMRFamide on trochozoan larvae and those focus on the regulation of the ciliary-based locomotion, which ultimately influences the vertical swimming direction [24-26]. In the veliger larvae of *Tritia obsoleta* it has been observed that FMRFamide induces twitching of the velum that results in ciliary arrests while contracted [24]. Many studies on adult trochozoans show diverse effects of FMRFamide on various muscles [33-36] and further taxon-specific functions such as osmoregulation [75], chromatophore expansion [76] or suppression of salivary gland activity [77]. Experiments on different adult bivalves showed a taxon specific up- or down regulation of the heartbeat by FMRFamide [32] and the experiments on trochozoan larvae showed a taxon specific up- or down regulation of the ciliary beating [24-26]. The various taxon specific effects and association with different tissues suggest that the FaRP receptor-ligand pair proved to be generally useful as a regulating signaling system and was likely redeployed several times during trochozoan evolution.

## Supporting material and data accession

Sequences of the here newly described genes and accession numbers of published sequences that were used in this study are listed in the supplementary material. Pictures of chromogenic *in situ* hybridization and a SEM picture of an early larva are shown in sFig.1. A list of tested peptides is given in sTab. 1.

## Acknowledgements

We want to thank Chema Martin Duran, Bruno Vellutini, Carmen Andrikou and Yale Passamaneck for helping with animal collection, spawning and fixation and Kevin Pang for reading the manuscript. The generous help of the “Centennial” crew and the administration of the Friday Harbor Laboratories, WA, USA made the animal collection and laboratory work possible. We further thank Jürgen Berger from the Max-Planck-Institute for Developmental Biology (Tübingen, Germany) for taking the beautiful SEM pictures of the larvae. This research was supported by the FP7-PEOPLE-2012-ITN grant no. 317172 “NEPTUNE” and received further support by the DFG - Deutsche Forschungsgemeinschaft to GJ (Reference no. JE 777/3-1).

## Supplementary Material

**sFig.1:**
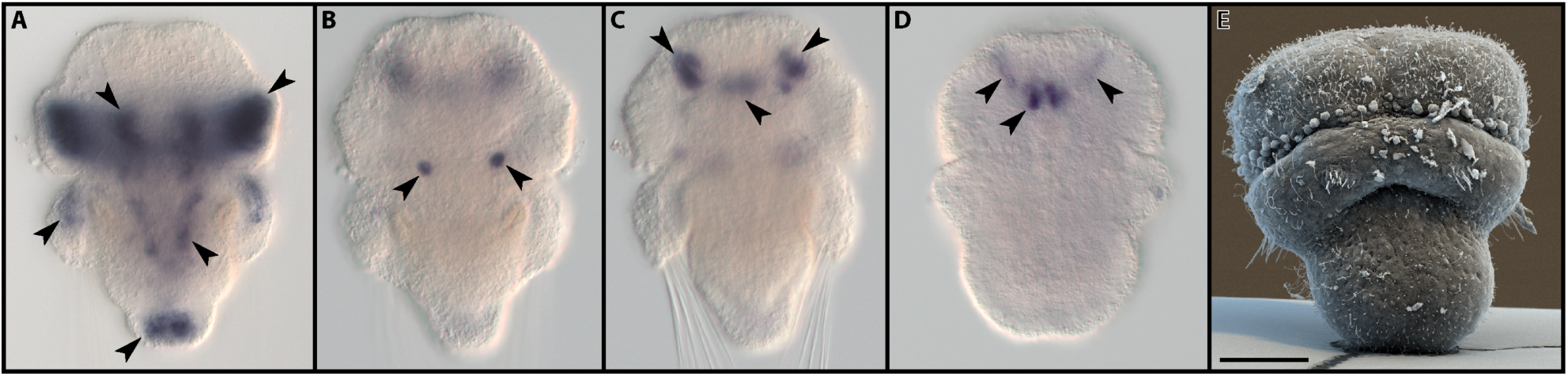
FLRFamide peptide and receptor expression in *T.transversa* larvae. **A-C** late larvae, *in situ* hybridization. Arrows indicate expression domains. **A** FLRFamide receptor expression. **B** ventral FLRFamide expression between apical and mantle lobe. (Same specimen as in C, but with a focus on the ventral side.) **C** FLRFamide expression in apical lobe. **D** early larva with FLRFamide expression in apical lobe. **E** SEM picture of early larva. Scale bar = 30 *μ*m.

**sTab.1:**
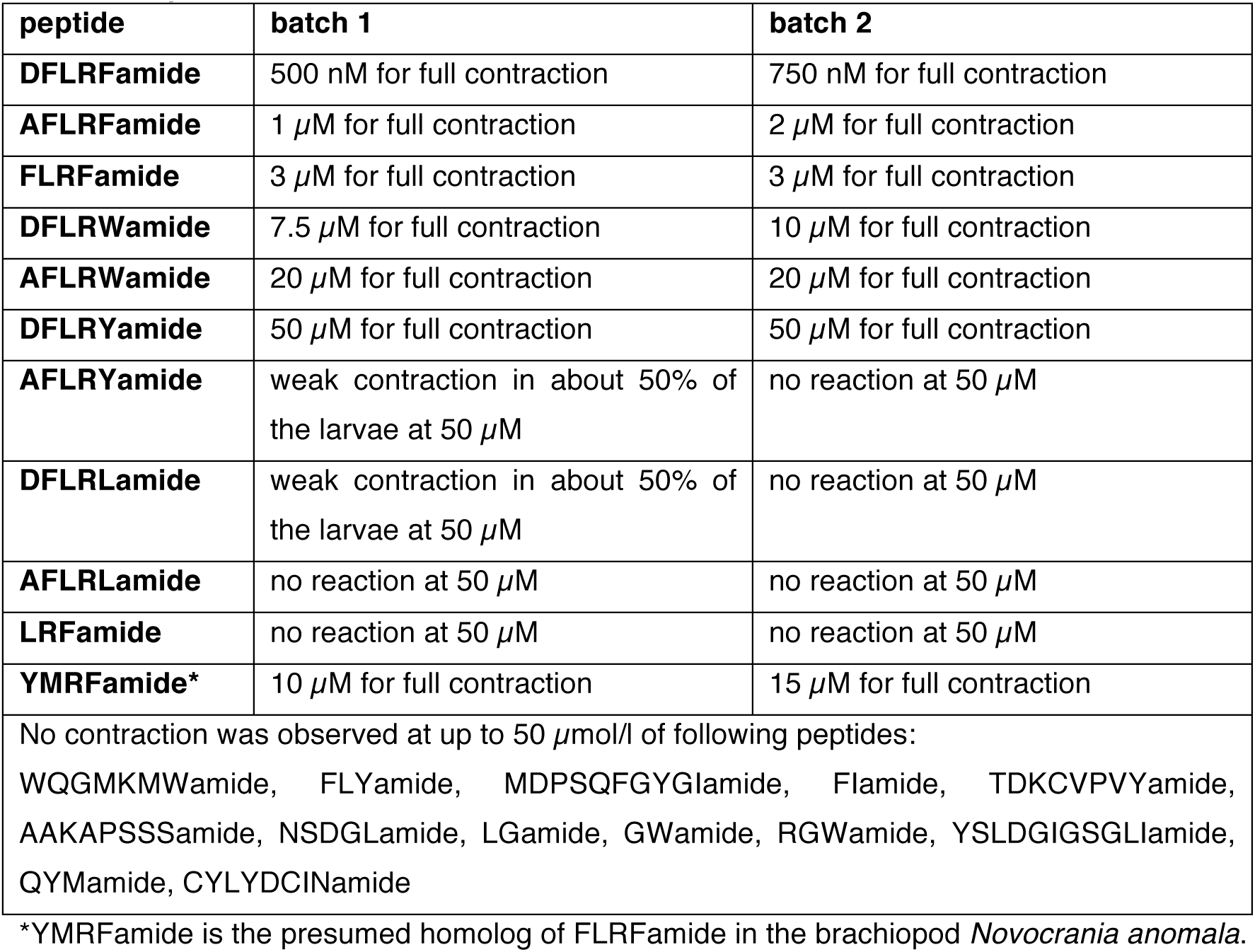
Peptide concentrations that lead to a contraction of the larvae.

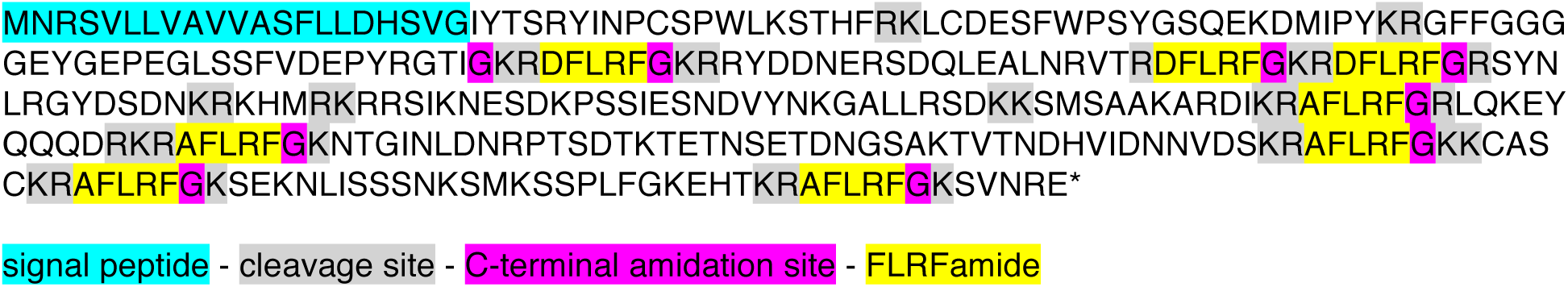
*Terebratalia transversa* FLRFamide prepropeptide sequence.

## FLRFamide precursor nucleotide sequence (Ttra.rna.tri.29781.1, completed with RACE PCR)

ATGAACAGGTCTGTGTTACTGGTTGCAGTCGTAGCTTCATTTTTGCTGGACCATTCAGTTGGAATCTATACTAGTCGCTATATTAATCCATGTTCGCCTTGGCTCAAATCTACCCATTTCAGAAAACTATGCGACGAGAGCTTCTGGCCATCCTACGGTAGCCAAGAAAAAGATATGATTCCATATAAGCGAGGTTTTTTTGGCGGTGGAGGAGAATATGGTGAACCAGAAGGACTGTCGAGCTTTGTTGATGAGCCGTATAGAGGAACAATAGGAAAACGAGATTTTCTGAGATTCGGAAAGAGACGATATGATGACAATGAACGAAGTGATCAATTGGAAGCACTAAATCGTGTCACCAGAGATTTTTTGAGATTTGGAAAACGAGATTTCTTAAGATTTGGCAGGAGCTATAATCTACGTGGATATGACTCAGACAACAAAAGAAACCATATGCGAAAGCGCCGTTCGATAAAGAATGAATCGGATAAGCCTAGTAGTATTGAGTCTAATGACGTTTACAACAAGGGAGCCCTTTTAAGGTCAGACAAGAAGTCAATGTCCGCCGCAAAAGCACGGGACATCAAACGAGCATTTCTAAGATTTGGTCGACTTCAAAAGGAATATCAGCGACAAGACCGAAAACGCGCATTTTTGCGTTTTGGAAAGAACACTGGAATAAATTTGGATAATAGGCCTACATCTGATACAAAAACTGAAACGAATTCAGAAACAGACAATGGTTCAGCGAAGACAGTAACAAATGATCACGTGATTGACAATAATGTTGACAGCAAACGCGCCTTTTTAAGATTTGGAAAAAAATGCGCAAGTTGTAAAAGAGCATTTTTACGTTTTGGGAAGTCTGAGAAGAATTTAATAAGTAGCTCGAACAAGTCGATGAAATCGAGCCCTCTCTTTGGGAAAGAACATACGAAACGAGCGTTTTTGAGGTTTGGAAAAAGTGTGAACAGAGAA

## Terebratalia transversa FLRFamide receptor (deorphanized)

MAAAKEFPGIIRKRESYFKPVYIVNDKLLLNTTTTIRATTGLPLGHDTTMVTSMELNCTLPNCTNTTTNDTTNGGSPIPLADSAQALIITFYTLAIILAAFGNILAIIIFSTGRRSRGDLRTYLLNLALADLAMSVFCIPFTFPTIIYHYWQFGSAMCTIVLFLQTATVVVSVSTNMAIGIDRFLAVTFPLRSRASRKQKVVKRIVVVVIWCLSFLLASPNFVVAQTVDLGNGYYQCTEKWPGGKQPKLIFGIFILIFTYIIPLVILLTTYGIIAMKLWRRQAPGEANRARDEAQLQSKRKVIKMLFTIVLLFGVCWLPLHTFINVLDFNPELAQNADQRILEIIYICFHWLAMSNSFQNPIIYGFLNDNFRADFWDLIFVCLPCCSKAKYFKNHRRMSYTRSPANRWQQSSFSHLERRASARPLRNGSTSSSSDTDKHNKFNSFARSKTDTALLGVHEPSKGRTFIKRNGAKRSKQPMALVVPTVTYNSRDNDVTSREMEKLLEGIEPDTSEKTNSIK*

## Terebratalia transversa orphan luqin GPCR homolog

FISMFSAITVLSITGNVLVCFAVLRNQTMRSSSYFFILNLAVSDILMATMCIPFTFVANVLLDWWPFGHVMCPLVNFLQAMAVFLSAFTLIVISLDRYVAIIFPLKARLTTKQIKVVIGIVWCCAVIPPIPIAVYGRVIRYVGRAYCQEIWTDNNNRFVYSLCILGLQFFVPLMVLVYTYIRIGIVIWGKKFPGEAEYNRDQRMATSKRKMVKMMMMVVIMYIICWLPYHCITIHGSIDDTFYDEEHAPTLWVFAYWLAMSNSCCNPLIYFYMNSKFRRSFKKTFCLLCCCKRISGRTCQESVKIKRTNTYATNTTTKESSG SKNGTTVKTSCSSRQRTSGDELAMIDILPV*

## Terebratalia transversa orphan neuropeptide GPCR, related to trochozoan NKY and insect FMRFa receptor

MTCLRTMLENTSPTGKLSMSREITNMTAGLTGVNQTYGNMSVCGHIPDPSTDIIMFQFIAWGIIGSILVLGGCVGNILAIIVLNRHSMGTFTSTYLSALAIFDTILLLCFLFSFSLPTIWNITWDSTYIDIIYPKMHLVIYPLTLISQQCTIYVTVAFTIQRYCAINWPLKRNKCLLSSRTQALIVITILILGSVIYNSPRMIEFTFYQCYSLQTNQVLQKIVPSEFGSDPTFRKVYHIYLFISVIFMVPFLVLVIFNTLLWLAVRRSKKLQIQKASTVKENNITIMLIAIVVVFLICQILPIADNIFMVTLSTATLNNNKYIKFTTISNLMVALNSSINFILYCMFGQRFRQIFLNLFCCKELNINFEGSRSIRWTRLSSFRSTLRDNKDGGNQPLRDNKDGGNQ

## Terebratalia transversa orphan neuropeptide GPCR

LRHLTPLLTLSLVGRAHFFILKSKIIYYWSCRIMSKELLSYLERFNGDKNLTTPMGNADRTYSNIGIIIAYTVIIVISLFGNVLVCQVVYRNKRLQTVTNIFIVNLAIADILMSSLNIPFTITRLTLDEWVLGSFVCHVFYFIPMVSVYVSTFTLTLIAIDRHQVIVYPLRPKITKRYGVIVLGVIWTLAITLAAPFAILAREADIHIVSREVKWCKMDYPHPSVLFHDSITLITISIQYCLPFVIISIMYGRIAKRLWSRGPLGHLTRAQELHHSKTKKKSIRMLVIVVCIFGLCWLPLNLYHILTDFHTDKLTFRHNSKAFIACHWLAFSSVGYNPFVYCWLNDAFRKEVKHILQCSRKDDSKIHPRGADKKKQQPSLTTRRTSSIRSTYISSSKVKKDTDIQGKISPDYQGDIDMQDKISPDYQQLTAMLQQDVPMRDINQLTNANQYSERAYPKAMQRDSESISPDERSLIEALRGAPHASEEDLDDIL*

## Accession numbers of neuropeptide receptor reference sequences (fig. 3)

[Xboc.rna.tri.15475.1 Xenoturbella bocki, in transcriptome SRX1343818] [Nwes_Locus_45236.0_Transcript_1/0 Nemertoderma westbladi, in transcriptome SRX1343819] [Msti.rna.tri.15359.1 Meara stichopi, in transcriptome SRX1343814] [Msti.rna.tri.31113.1 Meara stichopi, in transcriptome SRX1343814] [Msti.rna.tri.31092.1 Meara stichopi, in transcriptome SRX1343814] [Hspi_Locus_51813.1_Transcript_3/0 Halicryptus spinulosa, in transcriptome SRX1343820] [AKQ63075.1 Platynereis dumerilii, Luqin receptor, deorphanized] [AKQ63063.1 Platynereis dumerilii, FMRFamide receptor, deorphanized] [O44426 Lymnaea stagnalis, Luqin receptor, deorphanized] [P92045 Lymnaea stagnalis, Lymnokinin receptor, deorphanized] [P49146 Homo sapiens, Neuropeptide Y receptor type 2, deorphanized] [P97295 Mus musculus, Neuropeptide Y receptor type 2, deorphanized] [Q9ERC0 Rattus norvegicus, Neuropeptide Y/peptide YY-Y2 receptor, deorphanized] [P29371 Homo sapiens, Neuromedin-K receptor, deorphanized] [P47937 Mus musculus, Neuromedin-K receptor, deorphanized] [P16177 Rattus norvegicus, Neuromedin-K receptor, deorphanized] [FBpp0076853 Drosophila melanogaster, Leucokinin receptor, deorphanized] [FBpp0084470 Drosophila melanogaster, RYamide receptor, deorphanized] [FBpp0081791 Drosophila melanogaster,Tachykinin receptor 1, deorphanized] [FBpp0084873 Drosophila melanogaster, Tachykinin receptor 2, deorphanized] [ELT88896 Capitella teleta] [XP_009016737 Helobdella robusta] [XP_009054576 Lottia gigantea] [XP_009060043 Lottia gigantea] [XP_005090267 Aplysia californica] [EKC27293 Crassostrea gigas] [XP_009027087 Helobdella robusta] [XP_007899584 Callorhinchus milii] [XP_009054574 Lottia gigantea] [BAO01094 Nilaparvata lugens] [XP_002730513 Saccoglossus kowalevskii] [XP_002596257 Branchiostoma floridae] [XP_002734699 Saccoglossus kowalevskii] [XP_002738788 Saccoglossus kowalevskii] [XP_002731479 Saccoglossus kowalevskii][XP_002742045 Saccoglossus kowalevskii] [XP_002596255 Branchiostoma floridae] [NP_001161681 Saccoglossus kowalevskii] [XP_002732003 Saccoglossus kowalevskii] [XP_006812800 Saccoglossus kowalevskii][XP_009060304 Lottia gigantea] [XP_003700723 Megachile rotundata] [XP_002732001 Saccoglossus kowalevskii] [NP_001161604 Saccoglossus kowalevskii] [NP_001098693.1Takifugu rubripes] [XP_001342488.2 Danio rerio] [XP_009064591.1 Lottia gigantea] [XP_009067028.1 Lottia gigantea] [XP_009050865.1 Lottia gigantea] [XP_009064514.1 Lottia gigantea] [ELT99672.1 Capitella teleta] [XP_009017792.1 Helobdella robusta] [XP_009017796.1 Helobdella robusta] [XP_009062052.1Lottia gigantea] [XP_008498708 Calypte anna] [Q1ACB1 Oncorhynchus mykiss] [F1R5V3_DANRE Danio rerio] [E9HAW0_DAPPU Daphnia pulex] [G3X054_SARHA Sarcophilus harrisii] [G1NS97_MELGA Meleagris gallopavo] [J9JKR1_ACYPI Acyrthosiphon pisum][B3XXN5_BOMMO Bombyx mori] [B4MM03_DROWI Drosophila willistoni] [K1PQW2_CRAGI Crassostrea gigas] [B3XXN2_BOMMO Bombyx mori] [H3ILX9_STRPU Strongylocentrotus purpuratus] [H3ILY0_STRPU Strongylocentrotus purpuratus] [H9K8U7_APIME Apis mellifera] [Q8VHD7_RAT Rattus norvegicus] [G1TPU6_RABIT Oryctolagus cuniculus] [K1Q8V2_CRAGI Crassostrea gigas] [D6WD17_TRICA Tribolium castaneum][Q6AWE5_DROME Drosophila melanogaster] [I4IY86_TAKRU Takifugu rubripes] [F1R3V0_DANRE Danio rerio] [F7E6B1_MONDO Monodelphis domestica] [B4JUW2_DROGR Drosophila grimshawi] [Q94736_STOCA Stomoxys calcitrans] [E9FUQ7_DAPPU Daphnia pulex] [Q8T8D1_UREUN Tachykinin receptor]

